# Transcription initiation at a consensus bacterial promoter proceeds via a “bind-unwind-load-and-lock” mechanism

**DOI:** 10.1101/2021.03.28.437135

**Authors:** Abhishek Mazumder, Richard H Ebright, Achillefs N Kapanidis

## Abstract

Transcription initiation starts with unwinding of promoter DNA by RNA polymerase (RNAP) to form a catalytically competent RNAP-promoter complex (RP_O_). Despite extensive study, the mechanism of promoter unwinding has remained unclear, in part due to the transient nature of intermediates on path to RPo. Here, using single-molecule unwinding-induced fluorescence enhancement to monitor promoter unwinding, and single-molecule fluorescence resonance energy transfer to monitor RNAP clamp conformation, we analyze RPo formation at a consensus bacterial core promoter. We find that the RNAP clamp is closed during promoter binding, remains closed during promoter unwinding, and then closes further, locking the unwound DNA in the RNAP active-centre cleft. Our work defines a new, “bind-unwind-load-and-lock,” model for the series of conformational changes occurring during promoter unwinding at a consensus bacterial promoter and provides the tools needed to examine the process in other organisms and at other promoters.

**Significance statement:** Transcription initiation, the first step and most important step in gene expression for all organisms, involves unwinding of promoter DNA by RNA polymerase (RNAP) to form an open complex (RPo); this step also underpins transcriptional regulation and serves as an antibiotic target. Despite decades of research, the mechanism of promoter DNA unwinding has remained unresolved. Here, we solve this puzzle by using single-molecule fluorescence to directly monitor conformational changes in the promoter DNA and RNAP in real time during RPo formation. We show that RPo forms via a “*bind-unwind-load-and-lock*” mechanism, where the promoter unwinds outside the RNAP cleft, the unwound template DNA loads into the cleft, and RNAP “locks” the template DNA in place by closing the RNAP clamp module.

## Introduction

Transcription initiation is the first and most highly regulated step in gene expression (*1,2*). During transcription initiation, RNA polymerase (RNAP), together with the transcription initiation factor σ, unwinds ∼13 bp of promoter DNA to form a “transcription bubble,” and places the template-strand ssDNA of the unwound transcription bubble in contact with the RNAP active center, yielding a catalytically competent RNAP-promoter transcription-initiation complex (RPo; *1*). High-resolution structures of RP_O_ define the contacts that RNAP and σ make with promoter DNA, as well as the conformation and interactions of unwound template-strand ssDNA engaging the RNAP active center (*3-5*). Structural and biochemical experiments suggest that transcription-bubble formation is initiated by unwinding the DNA base pair at the upstream end of the transcription bubble--breaking the base pair, and unstacking and flipping the non-template-strand base of the broken base pair, and inserting the unstacked and flipped non-template strand base into a protein pocket of σ --followed by propagation of the unwinding in a downstream direction (*1,2,5,6*). However, the mechanism by which DNA is unwound and loaded into the RNAP active-center cleft has remained controversial (reviewed in *1*). In crystal structures of RNAP-σ holoenzyme, the RNAP active-center cleft is too narrow to accommodate double-stranded DNA (<20 Å), and σ obstructs access of double-stranded DNA to the RNAP active-center cleft (*7-10*). As a result, there is no unobstructed path by which double-stranded DNA can access the RNAP active-center cleft in RNAP holoenzyme (*7-12*). Accordingly, it has remained unclear where, when, and how promoter DNA is unwound and loaded into the RNAP active-center cleft.

To address these questions, two classes of models have been proposed. One class of models, termed “*load-unwind”* models, propose that: (i) the RNAP active-center cleft opens, through the swinging outward of one wall, termed the “clamp,” of the active-center cleft, allowing loading of promoter DNA into the active-center cleft as double-stranded DNA; (ii) promoter DNA unwinds inside the active-center cleft; and (iii) the active-center cleft closes, through the swinging inward of the clamp, during or after DNA unwinding (*1-2,7-9,11,13-14*). The other class of models, termed “*unwind-load”* models, proposes that: (i) promoter DNA unwinds outside the RNAP active-center cleft, and (ii) unwound promoter DNA loads into the active-center cleft as single-stranded DNA (*1,10*). Some versions of the unwind-load model postulate that opening and closing of the RNAP clamp is required for DNA unwinding and DNA loading to occur (*15,16*). Other versions of the unwind-load model postulate that opening and closing motions of the RNAP clamp is not required for DNA unwinding and DNA loading (*17*).

Some previous results have been interpreted as supporting *load-unwind* models, including results from static DNA footprinting of trapped putative on-pathway intermediates in formation of RPo, suggesting the presence of double-stranded DNA inside the active-center cleft (*18-21*), results from kinetic DNA footprinting suggesting the existence of on-pathway intermediates having double-stranded DNA inside the active-center cleft (*22*), fluorescence resonance energy transfer (FRET) results showing clamp opening and closing in RNAP and in trapped putative on-pathway intermediates in formation of RPo (*14,23-25*), functional correlations between inhibition of clamp opening and closing with inhibition of formation of RPo (*14,23-24*), and crystal and cryo-EM structures of trapped putative on-pathway intermediates in formation of RPo having double-stranded DNA inside an open active-center cleft (*26*).

Other previous results have been interpreted as supporting *unwind-load* models, including time-resolved footprinting experiments suggesting that promoter unwinding occurs outside the active-center cleft and precedes rate-limiting conformational changes in RNAP (*27*), and cryo-EM structures of trapped putative on-pathway intermediates in formation of RPo containing partly unwound DNA inside a closed active-center cleft (*17*).

However, the previous results either have relied on analysis of artificially trapped complexes that have not been firmly established to correspond to *bona fide* on-pathway intermediates, or have relied on analysis of ensemble kinetics for which the identities and orders of appearance of intermediates have not been firmly established (*14,18-27*). Moreover, the previous results have been complicated by differences in the species source of the RNAP analyzed, differences in the σ factors analyzed, and differences in the promoter sequences analyzed (*14,18-27*). Here, we report the use of single-molecule\ kinetic studies to define the pathway of DNA unwinding and DNA loading, without the assumptions that complicate analysis of artificially trapped complexes, and without the heterogeneity, population averaging, and time averaging that complicate analysis of ensemble kinetics. We used single-molecule promoter unwinding-induced fluorescence enhancement (smUIFE) to monitor DNA unwinding in solution in real time during formation of RPo, and we used single-molecule fluorescence resonance energy transfer (smFRET) to monitor opening and closing of the RNAP clamp in solution in real time during formation of RPo. In all experiments, we analyzed *Escherichia* coli RNAP σ^70^ holoenzyme at a consensus bacterial core promoter, comprising a consensus −35 element, a consensus −10 element, and a consensus −35/-10 spacer.

## Results

### Single-molecule unwinding-induced fluorescence enhancement (smUIFE)

Previous work indicates that a promoter derivative having the fluorescent probe Cy3 site-specifically incorporated in the transcription-bubble region exhibits a ∼2-fold increase in fluorescence emission intensity upon promoter unwinding during RPo formation and exhibits a ∼2-fold decrease in fluorescence emission intensity upon promoter rewinding during promoter escape (*15,28*). These changes in fluorescence emission intensity provide a powerful approach to monitor promoter unwinding and rewinding in solution (*15,28*). Here, we have adapted this approach to enable detection of promoter unwinding in solution at the single-molecule level, and we designate our adaptation of the approach “single-molecule unwinding-induced fluorescence enhancement” (smUIFE), to underscore the similarity to the established method of single-molecule protein-induced fluorescence enhancement (smPIFE; *29*; Fig. 1A).

**Figure 1.**
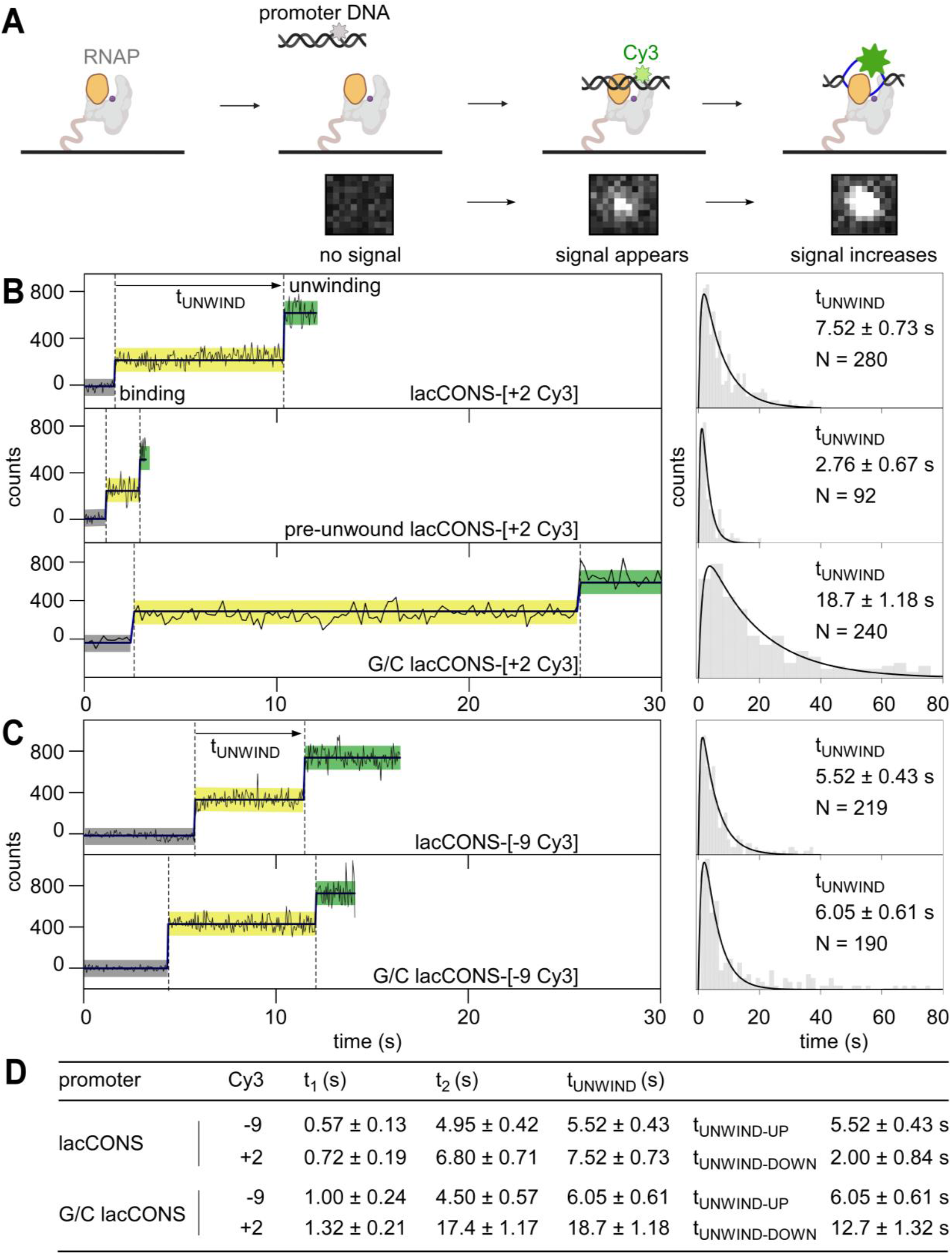
smUIFE: DNA unwinding in the upstream part of the transcription bubble precedes DNA unwinding in the downstream part of the transcription bubble. **A**. (*top*) Design of experiment monitoring promoter unwinding in real time. Grey, RNAP; orange, RNAP clamp; purple dot, RNAP active-center; black, ds-DNA; blue, ss-DNA; light green, Cy3 on ds-DNA; dark green, Cy3 on ss-DNA. (*bottom*) A cropped area (0.94 μm × 1.034 μm) of the field of view, showing appearance and enhancement of fluorescence signal from binding of single Cy3-labelled promoter fragment to an immobilised RNAP molecule. **B**. (*left*) Time-trajectories of intensity from Cy3 on downstream segment of promoter bubble. Black, raw intensity; dark blue, idealised intensity; Hidden Markov Model (HMM)-assigned states: no promoter (black bars), closed promoter (light yellow bars) and open promoter (green bars). Frame rates: 50-ms, top and middle; 200-ms, bottom. Laser powers: 0.60 mW, top and middle; 0.15 mW, bottom. (*right*) Dwell-time histograms of promoter state before unwinding, tUNWIND. **C**. (*left*) Time-trajectories of intensity from Cy3 on upstream segment of promoter bubble. Colors as in B. Frame rates: 50 ms. Laser powers: 0.6 mW. (*right*) Dwell-time histograms of promoter state before unwinding, tUNWIND. **D**. Table comparing unwinding times for different promoter constructs.

First, we analyzed a promoter DNA fragment having Cy3 incorporated at a site at the downstream edge of the transcription bubble (non-template strand position +2) of a consensus σ^70^-dependent bacterial promoter (lacCONS-[+2 Cy3]; Fig. S1A). Upon adding the Cy3-containing promoter DNA fragment to *E. coli* RNAP-σ^70^ holoenzyme immobilised on a coverslip mounted in a total-internal-reflection fluorescence (TIRF) microscope, we detected the appearance of fluorescence signal from single fluorescent species, indicating binding of single molecules of Cy3-containing promoter DNA fragment to surface-immobilised single molecules of RNAP holoenzyme (Fig. 1A, bottom). Control experiments confirmed that the Cy3-containing promoter DNA fragments were bound exclusively to immobilised RNAP molecules and showed that the majority (∼60%) of the resulting RNAP-promoter complexes were resistant to challenge with heparin, indicating they represented stable, sequence-specific complexes (Fig. S2A-B).

We extracted intensity-vs-time-trajectories for the formation of RNAP-promoter complexes, and identified different classes of trajectories. A large class of trajectories (∼45%) was characterised by the appearance of a fluorescence intensity of ∼200 counts, followed by an increase in fluorescence intensity to ∼450 counts, followed by either a decrease of the intensity to ∼200 counts or a disappearance of the intensity (Fig. 1B, top left; Fig. S3, *middle*). We assigned the states with no intensity as species that lack promoter DNA or that have promoter DNA with photobleached Cy3; the states with an intensity level of ∼200 counts as RNAP-promoter complexes having double-stranded DNA at the Cy3 incorporation site; and states with an intensity level ∼450 counts as RNAP-promoter complexes having single-stranded DNA at the Cy3 incorporation site. This assignment yields a reaction sequence comprising: binding of double-stranded promoter DNA to RNAP (∼200 counts; “pre-unwinding state”), followed by promoter unwinding (∼450 counts; “unwound state”), followed by promoter rewinding (∼200 counts) or probe photobleaching (∼0 counts).

Focussing on the class of trajectories that showed promoter unwinding after binding (as opposed to other classes of trajectories that did not show promoter unwinding after binding; Fig. S3 and *SI Methods*), we analysed trajectories further to define the kinetics of promoter unwinding (see *SI Methods*). Using Hidden Markov Modelling (HMM; *30*), we extracted dwell times for the pre-unwinding states and plotted a dwell-time histogram (Fig. 1B, *top right*). The dwell-time histogram showed a peaked distribution, indicating a non-Markovian process having more than one rate-limiting steps before the formation of the unwound state (*31*). After fitting this histogram to a two-exponential function (see SI *Methods; 31*), we estimated the average time spent in the pre-unwinding state to be ∼7.5 s (Fig. 1B, *top right*; Fig. 1D).

To assess whether the assay reports accurately on the kinetics of promoter unwinding, we analyzed altered promoter derivatives predicted to unwind *more quickly* or *more slowly* than lac*CONS*. To accelerate unwinding, we lowered the energy barrier for unwinding by introducing a non-complementary sequence at positions −10 to −4 relative to the transcription start site (pre-unwound lacCONS-[+2 Cy3]; Fig. S1). Time-trajectories for pre-unwound lacCONS-[+2 Cy3] were qualitatively similar to those for lacCONS-[+2 Cy3] in terms of the intensity increase (Figs. 1B, *middle left*, S4A) and the shape of the dwell-time histogram (Fig. 1B, *middle right*); however, the trajectories showed significantly shorter dwell times in the pre-unwinding state (∼2.8 s vs. ∼7.5 s; Fig. 1B, *middle right* vs. *top right*). To decelerate the process, we raised the energy barrier for unwinding by introducing a G/C-rich sequence at positions −4 to +1 relative to the transcription start site (G/C lacCONS-[+2 Cy3]; Fig. S1). Time-trajectories for G/C lacCONS-[+2 Cy3] showed significantly longer dwell times in the pre-unwinding state (∼18.8 s vs. 7.5 s; Fig. 1B, *bottom right* vs. *top right*; Fig. 1D). Taken together, these results show that dwell times in the pre-unwinding state depend on energy barriers for promoter unwinding, consistent with expectation that the dwell times report on the kinetics of promoter unwinding.

Next, to determine whether promoter unwinding occurs in one step, or in more than one step, we assessed whether unwinding of the upstream half of the transcription bubble (positions −11 to −5) coincides with, or does not coincide with, unwinding of the downstream half of the transcription bubble (positions −4 to +2). To probe unwinding of the upstream half of *GC* promoter bubble, we performed analogous smUIFE experiments using promoter derivatives having Cy3 incorporated in the upstream half of the transcription bubble, at template-strand position −9 (lacCONS-[-9 Cy3] and G/C lacCONS-[-9 Cy3];and Fig. S1). The resulting trajectories were similar in terms of intensity increases (Fig. S4B) and dwell-time-distribution shapes to those obtained with promoter DNA fragments having Cy3 incorporated in the downstream edge of the transcription bubble, but the dwell times in the pre-unwinding state were significantly shorter: ∼5.5 s vs. ∼7.5 s for lacCONS-[-9 Cy3] vs. lacCONS-[+2 Cy3] (Fig. 1C, *top right* vs. Fig. 1B, *top right*; Fig. 1D) and ∼6.0 s vs. ∼18.8 s for G/C lacCONS-[-9 Cy3] vs. G/C lacCONS-[+2 Cy3] (Fig. 1C, *bottom right* vs. Fig. 1B, *bottom right*; 1D). We conclude that, for the promoters analyzed, unwinding of the upstream half of the transcription bubble occurs faster than unwinding of the downstream half of the transcription bubble, and we conclude that we can estimate from our data both the reaction time required for unwinding of the upstream half of the transcription bubble, tUNWIND-UP, (from the lifetime of the pre-unwinding state when Cy3 is incorporated at position −9; ∼5.5 s for lac-CONS), and the reaction time required for the subsequent unwinding of the downstream half of the transcription bubble, tUNWIND-DOWN, (from the difference in lifetimes between the pre-unwinding state when Cy3 is incorporated at position +2 and the pre-unwinding state when Cy3 is incorporated at position −9; ∼2.0 s for lacCONS; Fig. 1D). Our results confirm previous results (*16,17,27, 32-35*), indicating that promoter unwinding proceeds in a step-wise fashion, in an upstream-downstream direction, and suggest that the step for upstream unwinding is slower than downstream unwinding for lacCONS (∼5.5 s vs. ∼2.0 s), whereas the step for downstream unwinding is slower than upstream unwinding for G/C lacCONS (∼6.0 s vs. ∼13.0 s).

### Single-molecule unwinding-induced fluorescence enhancement (smUIFE) in the presence of an inhibitor that prevents RNAP clamp opening

To determine whether promoter unwinding is affected by preventing opening of the RNAP clamp, we repeated our experiments in presence of myxopyronin (Myx; *14, 15, 24, 36-37*), an RNAP inhibitor that prevents RNAP clamp opening (*14, 24, 36;* state with E* ∼0.36 in Fig. S5D) and that allows formation of heparin-sensitive RNAP-promoter complexes but prevents formation of heparin-resistant RNAP-promoter complexes (*36*; Figs. S5A-B). We reasoned that, if clamp opening is obligatory for promoter unwinding, preventing clamp opening by addition of Myx should either prevent or delay downstream unwinding. We first performed smUIFE experiments in the presence of Myx using promoter DNA fragment lacCONS-[+2 Cy3], which has Cy3 incorporated at the downstream edge of the transcription bubble. The results showed a fluorescence intensity enhancement of ∼2.4-fold, consistent with unwinding of the downstream edge of the transcription bubble (Fig. 2B, *bottom;* Fig. S5C; see *15*) and showed a dwell time between initial binding and intensity enhancement of ∼4.1 s (Fig. 2C, *bottom*). We next performed smUIFE experiments in the presence of Myx using promoter DNA fragment lacCONS-[-9 Cy3], which has Cy3 incorporated in the upstream half of the transcription bubble. The results were essentially identical: a fluorescence intensity enhancement of ∼2.4 fold (Fig. 2B, *top*; Fig. S5C, *top*) and a dwell time between initial binding and intensity enhancement of ∼4.0 s (Fig. 2C). For both promoter derivatives, the observed fluorescence intensity enhancements were similar in the absence and presence of Myx (Figs. S4A-B, *top panels* vs. Fig. S5C), and the observed dwell times between initial binding and intensity enhancement times were shorter--not longer--in the presence of Myx than in the absence of Myx (∼3.5 s shorter for transcription-bubble downstream edge and ∼1.5 s shorter for transcription-bubble upstream half; Figs. 2B-C vs. Figs. 1B-C, *top panels*). We infer--contrary to the models in which clamp opening is obligatory for promoter unwinding--that preventing RNAP clamp opening does not prevent promoter unwinding, and increases, not decreases, the kinetics of promoter unwinding.

**Figure 2.**
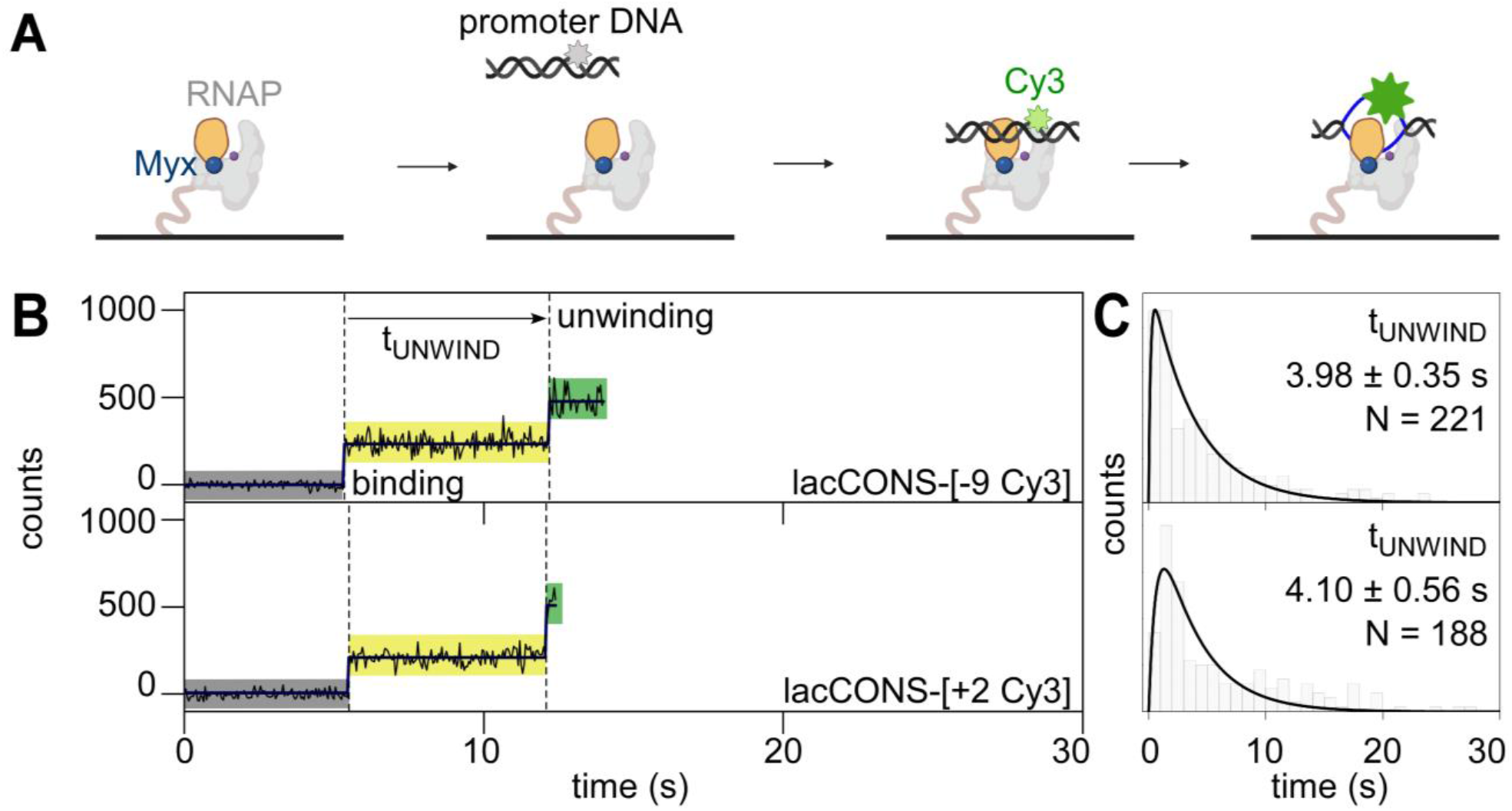
smUIFE in the presence of an inhibitor that prevents RNAP clamp opening: preventing RNAP clamp opening does not prevent DNA unwinding. **A**. Design of promoter unwinding experiment in presence of Myxopyronin (Myx). Blue sphere, Myx; rest as in 1A. **B**. (*left*) Time-trajectory of intensity from Cy3 on upstream (*top*) and downstream (*bottom*) segment of promoter bubble. Colors as in 1B. Frame rates: 50-ms. Laser powers: 0.60 mW. (*right*) Dwell-time histograms of promoter state before unwinding, tUNWIND.

### Single-molecule fluorescence resonance energy transfer (smFRET)

To assess *directly* whether RNAP clamp motions occur during RPo formation, we performed single-molecule fluorescence resonance energy transfer (smFRET) measurements using an RNAP derivative containing Cy3B, serving as a fluorescence donor, incorporated at the tip of the RNAP clamp and Alexa647, serving as a fluorescence acceptor, incorporated at the tip of the opposite wall of the RNAP active-center cleft, monitoring smFRET efficiency (E*) during RPo formation (Fig. 3A). In previous smFRET studies using the same probes, we showed that the RNAP clamp interconverts between open (E* ∼ 0.2), partly-closed (E* ∼ 0.3), and closed (E* ∼0.4) conformations in solution (*14, 23-25*). To monitor RNAP clamp motions during the formation of RP_O_, we immobilised promoter DNA molecules (biotin-lacCONS; Figs. S1, S6) on a coverslip mounted on a TIRF microscope, started recording, and added the doubly-labelled RNAP (Fig. 3A). Binding of doubly-labelled RNAP molecules to DNA was monitored by detecting the simultaneous appearance of donor and acceptor fluorescence emission signals on the surface, and RNAP clamp motions in RNAP-promoter complexes at and after binding were monitored by quantifying E* (Fig. S7A). Control experiments confirmed that the doubly-labelled RNAP molecules bound exclusively to immobilised promoter DNA molecules (Fig. S6A) and that the large majority (∼87%) of the complexes were resistant to heparin-challenge, indicating formation of stable, sequence-specific RNAP-promoter complexes (Fig. S6B).

**Figure 3.**
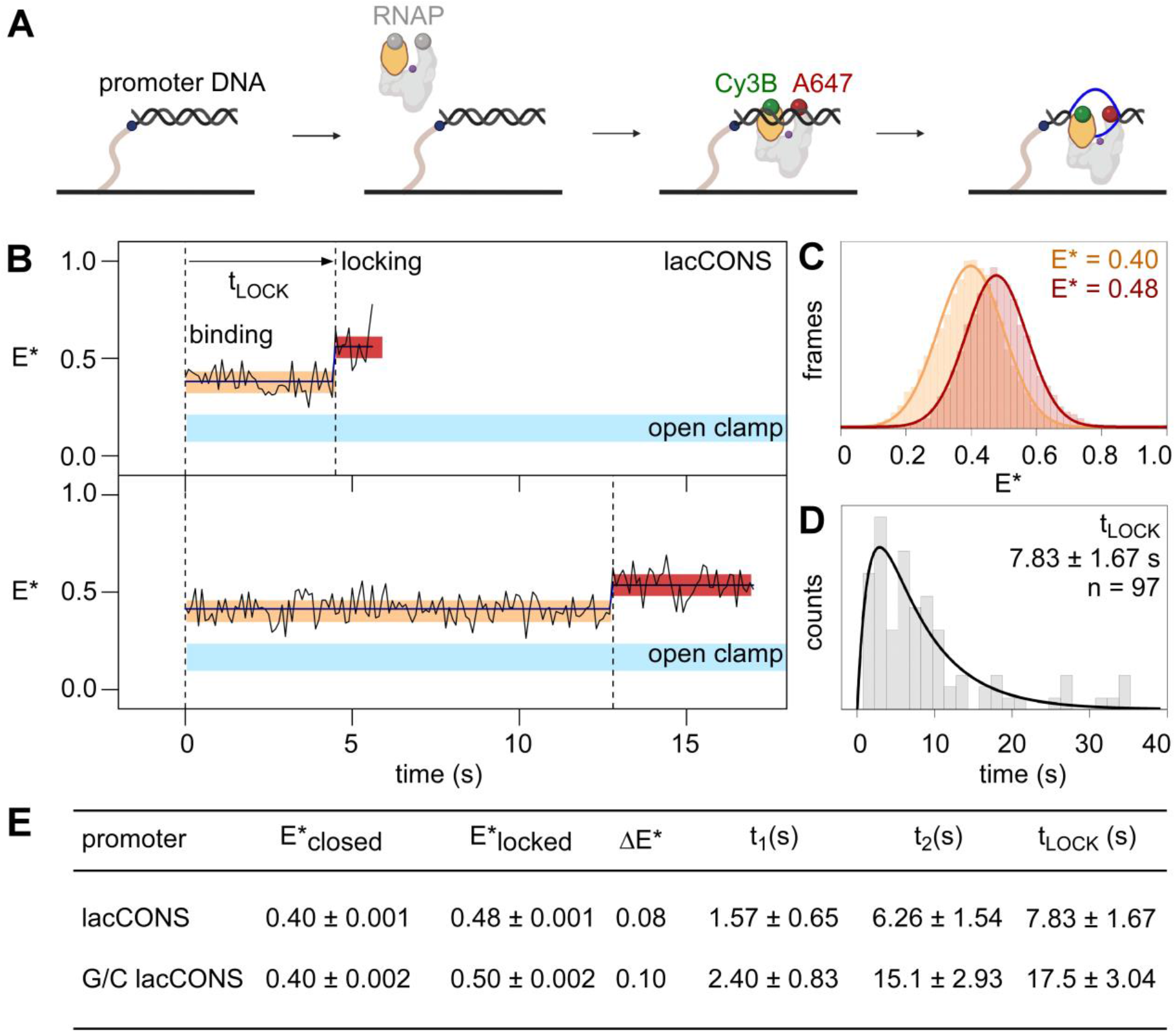
smFRET: DNA unwinding occurs without RNAP clamp opening and is followed by RNAP clamp locking. **A**. Design of experiment monitoring clamp status in real time. Black, ds-DNA; orange, RNAP clamp; grey, rest of RNAP; purple dot, RNAP active-center; blue, ss-DNA; green, Cy3B; and red, Alexa647. **B**. Representative time-trajectories of E* for experiments with a lacCONS promoter fragment, showing HMM-assigned closed clamp state (orange), locked-clamp state (red) and interstate transition (dark-blue). Expected range of E* values for an open clamp state is highlighted in light blue. Frame rate: 100-ms. Laser powers: 200 μW in red and 500 μW in green. **C**. HMM-assigned histograms and Gaussian fits of E* for full time-trajectories from experiments with a lacCONS promoter fragment. **D**. Dwell-time histograms of time before transition to the locked-clamp state, tLOCK, for experiments with a lacCONS promoter fragment. **E**. Table showing mean E*; difference in E* (ΔE*) between closed-clamp or locked-clamp states and time to transition to a locked-clamp state after initial binding for the lacCONS and lacCONS-GC promoter fragments.

In order to determine the RNAP clamp conformations immediately upon initial binding of RNAP to promoter DNA, we plotted distributions of E* values for the first five frames (0.5 s) after initial binding. The resulting distributions could be fitted to a Gaussian function with mean E* of ∼0.4, indicating that the initial binding of RNAP to promoter DNA involved RNAP with a closed clamp (Fig. S7B). Next, we examined full E* time-trajectories, for up to ∼60 s, following initial binding, seeking E* changes potentially consistent with RNAP clamp opening. We detected no E* changes--not even transient E* changes, within the ∼100 ms temporal resolution of the analysis--that potentially could be assigned as consistent with clamp opening (E* of ∼0.2; Fig. 3B, area highlighted in cyan; Fig. S7A).

Next, we examined full E* time-trajectories, following initial binding, seeking any E* changes potentially consistent with any RNAP clamp motions (see *SI Methods* and Fig. S8 for full classification). Intriguingly, a large fraction of trajectories (∼44%) started at E*∼0.40, indicative of the previously defined closed clamp state, then transitioned to E*∼0.48, indicative of a new, more tightly closed, clamp state, and then returned to E*∼0.40 or photobleached (Figs. 3B-C; Fig. S8B, *top*). We refer to the new, more tightly closed, clamp state with E*∼0.48, as the “locked-clamp” state. We further analysed E* time-trajectories to determine the time between initial binding of RNAP to promoter DNA and appearance of the locked-clamp state. The corresponding dwell-time histogram showed a peaked distribution and was fitted to a two-exponential function, yielding ∼7.8 s, as the time between initial binding of RNAP to promoter DNA and appearance of the locked-clamp state (Fig. 3D), a time that, within error, is identical to the ∼7.5 s time between initial binding and unwinding of the downstream half of the transcription bubble.

Analogous experiments with a promoter derivative having G/C-rich sequence at positions −4 to +1 relative to the transcription start site, G/C lacCONS, yielded similar results: i.e., initial binding by RNAP with a closed-clamp state (E*∼0.41; Fig. S9A; S9B, *left*); no trajectories showing transitions --not even transient transitions, within the ∼400 ms temporal resolution of the analysis--to an open-clamp state during a period up to ∼200 s after initial binding; a large fraction of trajectories (∼48%), showing a transition to a locked-clamped state (E*∼0.50, Fig. S9A; S9B, *right*), and a time between initial binding and appearance of the locked-clamped state matching the time required for unwinding of the downstream half of the transcription bubble (Figs. 3E, S9C).

We conclude that the RNAP clamp is in a closed state upon promoter binding, that the RNAP clamp does not open--not even transiently, within the temporal resolution of our analysis--between promoter binding and promoter unwinding, and that the RNAP clamp closes further--”locks”--after promoter unwinding.

## Discussion

Taken together, our results indicate that RPo formation by *E. coli* RNAP-σ^70^ holoenzyme at a consensus bacterial core promoter proceeds through a “*bind-unwind-load-and-lock*” mechanism, in which the RNAP clamp is closed upon promoter binding, remains closed during unwinding of promoter DNA--which proceeds in an upstream-to-downstream direction--and then closes further, locking the unwound DNA in the RNAP active-centre cleft (Fig. 4). The finding that the RNAP clamp remains closed during promoter unwinding strongly suggests that promoter unwinding occurs, at least in part, outside the RNAP active-centre cleft (Fig. 4).

**Figure 4.**
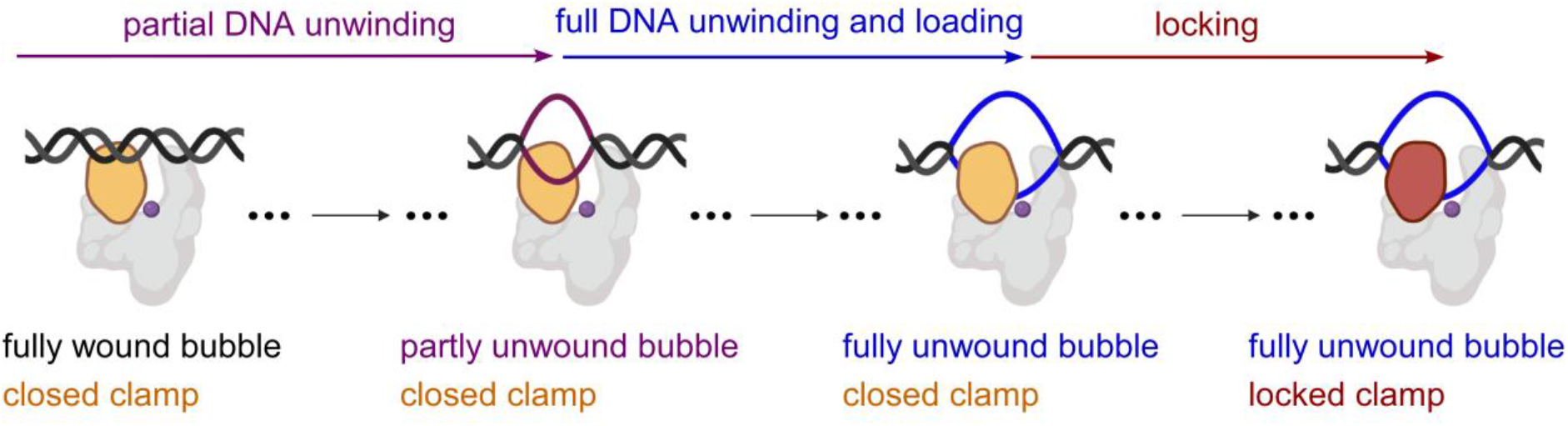
“Bind-unwind-load-and-lock” mechanism for the formation of RPo. Orange, closed clamp; red, locked clamp; grey, rest of RNAP; purple dot, RNAP active-center; black, ds-DNA; magenta, partly unwound bubble and blue, fully unwound bubble.

It remains to be determined whether a similar *bind-unwind-load-and-lock* mechanism pathway is used by *E. coli* RNAP σ^70^ holoenzyme at other promoters and by other *E. coli* RNAP holoenzymes. Consistent with the possibility that a similar mechanism is used by *E. coli* RNAP σ^70^ holoenzyme at other promoters, a recent cryo-EM study of transcription initiation by *E. coli* RNAP σ^70^ holoenzyme at another promoter, *rpsT* P2, identified structural states having a partly unwound transcription bubble and a closed RNAP clamp (*17*). We note that the smUIFE and smFRET methods reported in this work could be applied to analyze transcription initiation by any *E. coli* RNAP holoenzyme at any promoter and potentially could be adapted to analyze transcription initiation by any RNAP--bacterial, archaeal, or eukaryotic--at any promoter.

## Materials and Methods

Fluorescent probes were incorporated at specific sites in RNAP by use of unnatural-amino acid mutagenesis and Staudinger ligation (*23-25*). Full details of methods are presented in *[Supplementary methods](https://www.pnas.org/content/suppl/2020/06/20/1920427117.DCSupplemental)*.

## Data and software availability

Data necessary for replication are included in the submission. MATLAB software packages Twotone-ALEX and ebFRET are available, respectively, at https://kapanidis.web.ox.ac.uk/software and on GitHub (http://ebfret.github.io/).

## ACKNOWLEDGEMENTS

We thank Dr. Anssi Malinen for discussions and early work on the development of real-time smFRET assays, and Dr. Horst Steuer for the development of custom software. This work was supported by the Wellcome Trust [110164/Z/15/Z to A.N.K.] and NIH [GM041376 to R.H.E.].

## Author Contributions

A.M., R.H.E., and A.N.K conceived the project; A.M. prepared and characterized protein samples and performed single-molecule experiments; A.M. and A.N.K analyzed data; A.M., and A.N.K. prepared figures; and A.M., R.H.E., and A.N.K wrote the manuscript.

## Supplementary Methods

### Preparation of reagents: RNAP, Oligodeoxyribonucleotides, Myxopyronin

#### RNAP derivatives

For experiments in Figs. 1, 2, S2-S5, hexahistidine-tagged *Escherichia coli* RNAP holoenzyme was prepared using co-expression of genes encoding RNAP β’, β, α, ω and σ^70^ subunits to afford an RNAP σ^70^ holoenzyme derivative as follows: single colonies of *E. coli* strain BL21(DE3) (Millipore) co-transformed with plasmid pVS10 (*1*) and plasmid pRSFduet-sigma (*2*) were used to inoculate 20 ml LB broth (*3*) containing 100 μg/ml ampicillin and 50 μg/ml kanamycin and cultures were incubated 16 h at 37°C with shaking. Culture aliquots (2×10 ml) were used to inoculate LB broth (2×1 L) containing 100 μg/ml ampicillin and 50 μg/ml kanamycin; cultures were incubated at 37°C with shaking until OD_600_ = 0.6; IPTG was added to 1 mM; and cultures were further incubated 3.5 h at 37°C with shaking. Cells were harvested by centrifugation (4,000 x g; 20 min at 4°C), re-suspended in 20 ml buffer A (10 mM Tris-HCl, pH 7.9, 200 mM NaCl, and 5% glycerol), and lysed using an EmulsiFlex-C5 cell disrupter (Avestin). The lysate was cleared by centrifugation (20,000 x g; 30 min at 4°C), precipitated with polyethyleneimine (Sigma-Aldrich) as in (*4*), and precipitated with ammonium sulfate as in (*4*). The precipitate was dissolved in 30 ml buffer A and loaded onto a 5-ml column of Ni-NTA-agarose (Qiagen) pre-equilibrated in buffer A; the column was washed with 50 ml buffer A containing 10 mM imidazole, and eluted with 25 ml buffer A containing 200 mM imidazole. The sample was further purified by anion-exchange chromatography on Mono Q 10/100 GL (GE Healthcare; 160 ml linear gradient of 300-500 mM NaCl in 10 mM Tris-HCl, pH 7.9, 0.1 mM EDTA, and 5% glycerol; flow rate = 2 ml/min). Fractions containing hexahistidine-tagged *E. coli* RNAP σ^70^ holoenzyme were pooled, concentrated to ∼2 mg/ml using 30 kDa MWCO Amicon Ultra-15 centrifugal ultrafilters (EMD Millipore), and stored in aliquots at −80°C.

For experiments in Figs. 3 and S6-S9, fluorescently labelled, hexahistidine-tagged *E. coli* RNAP holoenzyme (hereafter ‘labeled-RNAP’) with Cy3B and Alexa647 at positions 284 on the β’ subunit, and 106 on the β subunit, respectively, was prepared using in-vivo reconstitution methods as described (*5*).

#### Nucleic acids

Oligodeoxyribonucleotides were purchased from IBA Lifesciences, dissolved in nuclease-free water (Ambion) to a final concentration of 100 μM and stored at −20°C. Oligodeoxyribonucleotides used in Figs. 1, S2-S5, were labelled with Cy3 N-hydroxysuccinimidyl (NHS) ester (Fisher Scientific) as described (*6*). Oligodeoxyribonucleotides were annealed by mixing two complementary strands in a ratio of 1:1 in hybridization buffer (50 mM Tris–HCl pH 8.0, 500 mM NaCl, 1 mM EDTA) and by heating for 5 min at 95°C, followed by cooling to 25°C in 2°C steps with 1 min per step using a thermal cycler (Applied Biosystems). Myxopyronin was prepared as described (*7*).

### Single-molecule fluorescence microscopy: smUIFE experiments

For experiments in Figs. 1, 2 and S2-S5, observation wells for real-time experiments were prepared as described (*5*). Briefly, a biotin-PEG-passivated glass surface was prepared, functionalized with Neutravidin (Sigma Aldrich), and treated with biotinylated anti-hexahistidine monoclonal antibody (Penta-His Biotin Conjugate; Qiagen), yielding wells with (biotinylated anti-hexahistidine monoclonal antibody)-Neutravidin-biotin-PEG-functionalised glass floors. Hexahistidine tagged RNAP σ^70^ holoenzyme molecules were immobilised in observation wells with (biotinylated anti-hexahistidine monoclonal antibody)-Neutravidin-biotin-PEG-functionalized glass floors, as follows: aliquots (30 μl) of 0.1 nM hexahistidine tagged RNAP σ^70^ holoenzyme in KG7 buffer (40 mM HEPES-NaOH, pH 7.0, 100 mM potassium glutamate, 10 mM MgCl_2_, 1 mM dithiothreitol, 100 μg/ml bovine serum albumin, and 5% glycerol) were added to the observation chamber and incubated 2-4 min at 22°C, solutions were removed, wells were washed with 2 x 30 μl KG7, and 30 μl KG7 imaging buffer (KG7 buffer containing 2 mM Trolox, 1 mg/ml glucose oxidase, 40 μg/ml catalase, and 1.4% w/v D-glucose) at 22°C was added.

For experiments in Fig. S2, Cy3-labelled promoter fragments were manually added (final concentration of 2 nM) to the observation wells containing immobilised RNAP σ^70^ holoenzyme molecules and incubated for 5 min; wells were then washed with KG7 and movies were recorded. Next, observation wells containing RNAP-promoter complexes were supplemented with KG7 containing 250 μg/ml heparin (Sigma Aldrich) solution, incubated for 1 min; wells were then washed with KG7 and movies were recorded.

For experiments monitoring RNAP-promoter complex formation reactions in real time (Figs. 1, 2, and S3-S5), observation wells containing immobilised RNAP were supplemented with KG7 imaging buffer, recordings were started, and Cy3-labelled promoter fragments were manually added (using a pipette) to the observation wells during the recording so as to yield a final concentration of 2 nM of Cy3-labelled promoter fragments in the wells.

For experiments in Fig. S5A-C, same procedures were followed, except that KG7 imaging buffer was supplemented with 20 μM Myxopyronin.

### Single-molecule fluorescence microscopy: smFRET experiments

For experiments in Fig. 3 and Figs. S6-S9, observation wells of biotin-PEG passivated glass surface was prepared and functionalized with Neutravidin (Sigma Aldrich) to yield Neutravidin-biotin-PEG-functionalised glass floors. Biotin-tagged promoter DNA fragments were then immobilised in these observation wells as follows: aliquots (30 μl) of 0.05 nM biotin-tagged promoter fragments in KG7 were added to the observation chamber and incubated 1 min at 22°C, solutions were removed, wells were washed with 2 x 30 μl KG7, and 30 μl KG7 imaging buffer at 22°C was added.

For experiments in Fig. S6, clamp-labelled RNAP molecules were manually added (final concentration of 2 nM) to the observation wells containing immobilised promoter DNA fragments, incubated for 5 min, wells were washed with KG7 and movies were recorded. Next, observation wells containing RNAP-promoter complexes were supplemented with KG7 containing 250 μg/ml heparin solution, incubated for 1 min, wells were washed with KG7 and movies were recorded.

For experiments monitoring RNAP-promoter complex formation reactions in real time (Figs. 3 and Figs. S7-S9, observation wells containing immobilised promoter fragments were supplemented with KG7 imaging buffer, recordings were started, and labeled-RNAP molecules (with Cy3B and Alexa647 at positions 284 on the β’ subunit, and 106 on the β subunit of RNAP σ^70^ holoenzyme) were manually added (using a pipette) to the observation wells during the recording so as to yield a final concentration of 2 nM of labeled RNAP in the wells.

### Single-molecule fluorescence microscopy: data collection

Single-molecule fluorescence experiments were performed using a custom-built objective-type total-internal-reflection fluorescence (TIRF) microscope (*10*). Light from a green laser (532 nm; Samba; Cobolt) and a red laser (635 nm; CUBE 635-30E, Coherent) was combined using a dichroic mirror coupled into a fiber-optic cable focused onto the rear focal plane of a 100x oil-immersion objective (numerical aperture 1.4; Olympus) and was displaced off the optical axis, such that the incident angle at the oil-glass interface of a stage-mounted observation chamber exceeded the critical angle, thereby creating an exponentially decaying evanescent wave (*11*). Alternating-laser excitation (ALEX; *8,9*) was implemented by directly modulating the green and red lasers using an acousto-optical modulator (1205C, Isomet). Fluorescence emission was collected from the objective, was separated from excitation light using a dichroic mirror (545 nm/650 nm, Semrock) and emission filters (545 nm LP, Chroma; and 633/25 nm notch filter, Semrock), was focused on a slit to crop the image, and then was spectrally separated (using a dichroic mirror; 630 nm DLRP, Omega) into donor and emission channels focused side-by-side onto an electron-multiplying charge-coupled device camera (EMCCD; iXon 897; Andor Technology). A motorized x/y-scanning stage (MS-2000; ASI) was used to control the sample position relative to the objective.

For experiments in Figs. 1-2, and Figs. S2-S5, frame rates were either 50-ms or 200-ms, and laser powers were either 0.60 mW or 0.15 mW at 532 nm. For experiments in Fig. 3 and Figs. S7-S8, frames were either 100-ms or 200-ms long, and laser powers were either 200 μ W in red, 500 μW in green, or 80 μW in red, 200 μW in green, respectively. For experiments in Fig. S9, frame rates were 400-ms long, and laser powers were 50 μ W in red, 150 μW in green. All data acquisition was carried out at 22°C.

### Single-molecule fluorescence microscopy: data analysis for smUIFE experiments

For experiments in Figs. 1-2, and Figs. S2-S5, localisation of single fluorescence emitters was detected and background-corrected fluorescence intensity-vs-time trajectories from each localisation were extracted and curated to exclude trajectories exhibiting very high fluorescence intensities (I_Cy3_) upon binding (I_Cy3_>750 photon counts), trajectories exhibiting multiple binding events during the observation window and trajectories exhibiting photoblinking. Initial inspection of time-trajectories for experiments with lacCONS-[+2 Cy3], revealed three main classes of molecules. Class-I molecules started with the appearance of signal having intensity of ∼200 photon counts, which remained stable until its disappearance (∼46%; S3, *top*). Class-II molecules also started with appearance of a signal having intensity of ∼200 photon counts, followed by intensity increase to ∼450 photon counts after some time, followed by either signal disappearance or intensity decrease to ∼200 photon counts (∼45%: Figs. 1B, S3, *middle*). Class-III molecules started with appearance of a signal having intensity of ∼450 photon counts, followed by intensity decrease to ∼220 photon counts, and subsequent signal disappearance, or intensity increase to ∼450 photon counts (∼9%; Fig. S3, *bottom*). Similar observations were made for all other Cy3-labelled promoters studied.

Based on intensity levels, we assigned states with no signal to absence of a promoter DNA or to a bleached probe; states with ∼200 photon counts to a promoter DNA that is double stranded (dsDNA) in the Cy3 vicinity; and states with ∼450 counts to a promoter that is unwound in the Cy3 vicinity. We thus assigned events in Class-I molecules to binding of a promoter DNA molecule to RNAP, followed by dissociation of complexes, indicating formation of non-specific complexes; these events are also consistent with bleaching occurring prior to any intensity increase. We assigned events in Class-II molecules to binding of a promoter DNA molecule to RNAP, followed by promoter unwinding, followed by bleaching or promoter rewinding. Finally, we assigned events in Class-III molecules to binding of an unwound promoter DNA molecule to RNAP, followed by promoter rewinding, followed by bleaching or promoter unwinding, indicating formation of complexes where the initial unwinding event is missed. We further curated intensity time-trajectories up to the point of initial ∼2-fold intensity enhancement for Class-II molecules only, since they showed an unambiguous bubble-opening event after initial binding.

Next, photon counts (I_Cy3_) for the set of curated intensity-vs-time trajectories were divided by 1000 to obtain I_Cy3_*; the corresponding I_Cy3_*-vs-time trajectories were analyzed to identify the different fluorescence intensity states using Hidden Markov Modelling (HMM) as implemented in the MATLAB (MathWorks) software package ebFRET (*12*) using a three-state model (one of which corresponds to a negligible I_Cy3_* state). The intensity I_Cy3_*, for each of the other two HMM-derived states were extracted from ebFRET and converted to I_Cy3_ values, which in turn were binned and plotted as I_Cy3_ count histograms (Fig. S4, S5C). These histograms were fitted using Gaussian distributions in Origin to define the mean fluorescence intensity that corresponds to each state (Fig. S4, S5C).

Dwell times for each fluorescence intensity states were extracted from HMM fits to I_Cy3_*-vs-time trajectories, were binned, and were plotted as distribution histograms in Origin. For experiments in Figs. 1-2, dwell-time distributions corresponding to the pre-unwinding state resembled peaked distributions indicating presence of at least two sub-steps and were fit to a two-exponential function of the form y = A*(e^-x/t1^ – e^-x/t2^), where y represents counts of dwells in the closed-bubble state following the initial binding event and before the signal increase event, x represents time, and t_1_ and t_2_ represent the lifetime of the individual sub-steps. From the fit, we estimate the lifetimes corresponding to the two sub-steps, as well as the total time spent in the first closed-bubble state (as the sum of time spent in the two sub-steps; Fig. 2B, right panels).

We note that it is possible that the transition involves more than two steps, but our current analysis and available data sets cannot specify the number and duration of each sub-step; consequently, we interpret qualititatively the shape of the dwell-time histogram as an indication of a multi-step process, and focus on the total time spent before the intensity enhancement event, since that can be interpreted with certainty as: the time taken before unwinding in the vicinity of Cy3.

### Single-molecule fluorescence microscopy: data analysis for smFRET experiments

For experiments in Fig. 3 and Figs. S6-S9, localizations in donor-emission (green) and acceptor-emission (red) channels were detected using the peak-finding algorithm of the MATLAB (MathWorks) software package Twotone, as described (*10*). Peaks detected in both emission channels (i.e., peaks for molecules containing both donor and acceptor probes) were fitted with two-dimensional Gaussian functions to extract background-corrected intensity-vs.-time trajectories for donor-emission intensity upon donor excitation (I_DD_), acceptor-emission intensity upon donor excitation (I_DA_), and acceptor-emission intensity upon acceptor excitation (I_AA_), as described (*10*). Intensity-vs.-time trajectories were curated to exclude trajectories exhibiting I_DD_ <100 or >1,000 counts or I_AA_ <200 or >1,000 counts, trajectories exhibiting multiple-step donor or acceptor photobleaching, trajectories exhibiting donor or acceptor photo-blinking and portions of trajectories following donor or acceptor photobleaching. Intensity-vs.-time trajectories were used to calculate trajectories of apparent donor-acceptor smFRET efficiency (E*) as described (*8-10*):

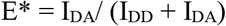

E* time trajectories were analyzed globally to identify E* states by use of HMM as implemented in MATLAB (MathWorks) software package ebFRET (*12*).

HMM analysis of E* time-trajectories revealed four types of molecules. The most abundant were Class-I molecules (∼45% of all events), that started with E*∼0.40, remained at E*∼0.40, and then bleached (Fig. S8A), and Class-II molecules (∼44%) that started with E*∼0.40, transitioned to E*∼0.48, and then bleached or returned to E*∼0.40 (Fig. S8B, *top*). More rarely, we observed Class-III molecules (∼9% of all events), that started with E*∼0.48, transitioned to E*∼0.40, followed by bleaching or transition to E*∼0.48; Fig. S8C), and, very rarely, we observed Class-IV molecules (∼2%), which started at E*∼0.40, switched between E*∼0.40-0.20 and bleached (Fig. S8D). We refer to the new E*∼0.48 state observed here as the “locked-clamp” conformation, with Class-II molecules providing information on the time taken to form the “locked-clamp” conformation; in contrast, Class-I molecules represent RNAP molecules that either bleach before the transition to the locked-clamp state or bind non-specifically to DNA; and Class-III molecules represent molecules where the initial clamp locking is missed.

E* time trajectories for Class-II molecules, were fitted to a two-state HMM model, E*-values from the fitted model were extracted, binned and plotted using Origin (Origin Lab), and were fitted to Gaussian distributions using Origin (Fig. 3B, right panels and Fig. S8B, left panel; colored curves). The resulting histograms provide population distributions of E* states and, for each E* state, define mean E* (Fig. 3B, right panels and Fig. S9B, right panel; colored bars and inset). Dwell-time distributions corresponding to time spent before the first transition from a closed-clamp state to a locked-clamp state were extracted from HMM fits to E*-vs-time trajectories, and were binned and plotted as distribution histograms in Origin (Fig. 3 and Fig. S9C). Dwell-time histograms obtained in this manner exhibited the shape of a peaked distribution, indicating presence of at least two sub-steps, were fitted to a two-exponential function of the form y = A*(e^-x/t1^ – e^-x/t2^), where y represents counts of dwells in the closed-clamp state following the initial binding event and before the clamp-locking event, x represents time, and t_1_ and t_2_ represent the lifetimes of the individual sub-steps. From the fit, we estimate the time spent in the two sub-steps, as well as the total time before transition to “locked-clamp” conformation (tLOCK) from the sum of times corresponding to the two sub-steps. Similar to the case for fluorescence enhancement experiments described previously, we note that there may be more than two-steps which contribute significantly to these dwell-times, and we could not infer an accurate model in terms of the number and duration of each sub-step involved from this dataset. Therefore, we avoid assigning the two sub-steps to specific conformations or events and focus on the total time spent before the transition to “locked-clamp” conformation.

## Supplementary Figures

**Fig. S1.**
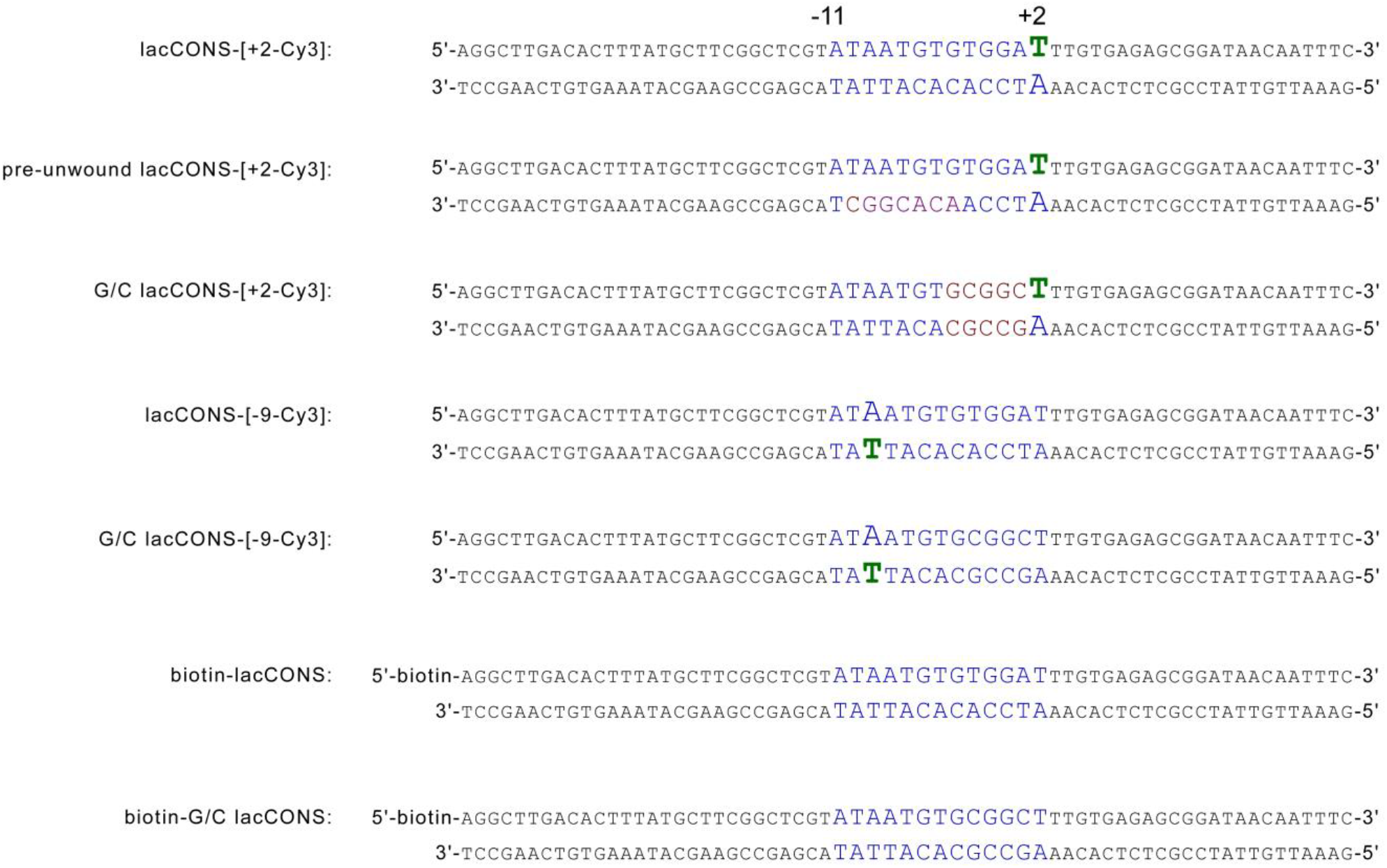
Sequence of consensus lac-promoter fragments used in the study. Top strand: non-template DNA strand; bottom strand: template strand. Nucleotides labelled with Cy3 are shown in green. The numbering refers to the DNA position relative to the transcription start site.

**Fig. S2.**
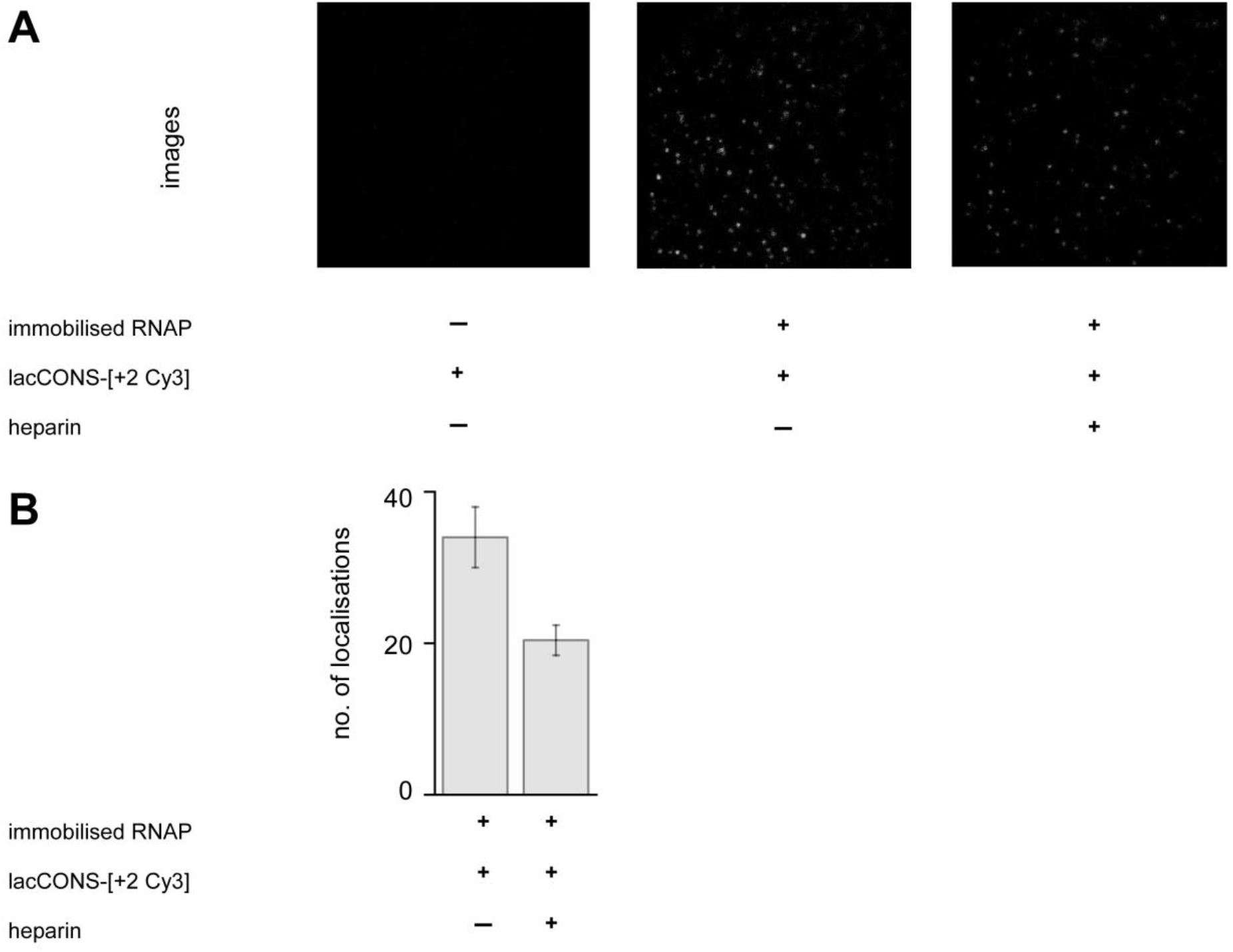
Characterisation of complexes formed between immobilised RNAP and Cy3-labelled promoter fragments. A. Raw images showing field of view (24 μm × 24 μm) with only immobilised hexahistidine-tagged RNAP holoenzyme (*left*), immobilised hexahistidine-tagged RNAP holoenzyme bound to Cy3-labelled promoter fragment (*middle*), and remaining immobilised hexahistidine-tagged RNAP holoenzyme complexes bound to Cy3-labelled promoter fragment after addition of heparin (*right*). **B**. Mean number of localisations per field of view for single Cy3-labelled promoter fragments bound to immobilised hexahistidine-tagged RNAP holoenzyme in absence and presence of heparin. Errors bars represent the standard deviation from the mean.

**Fig. S3.**
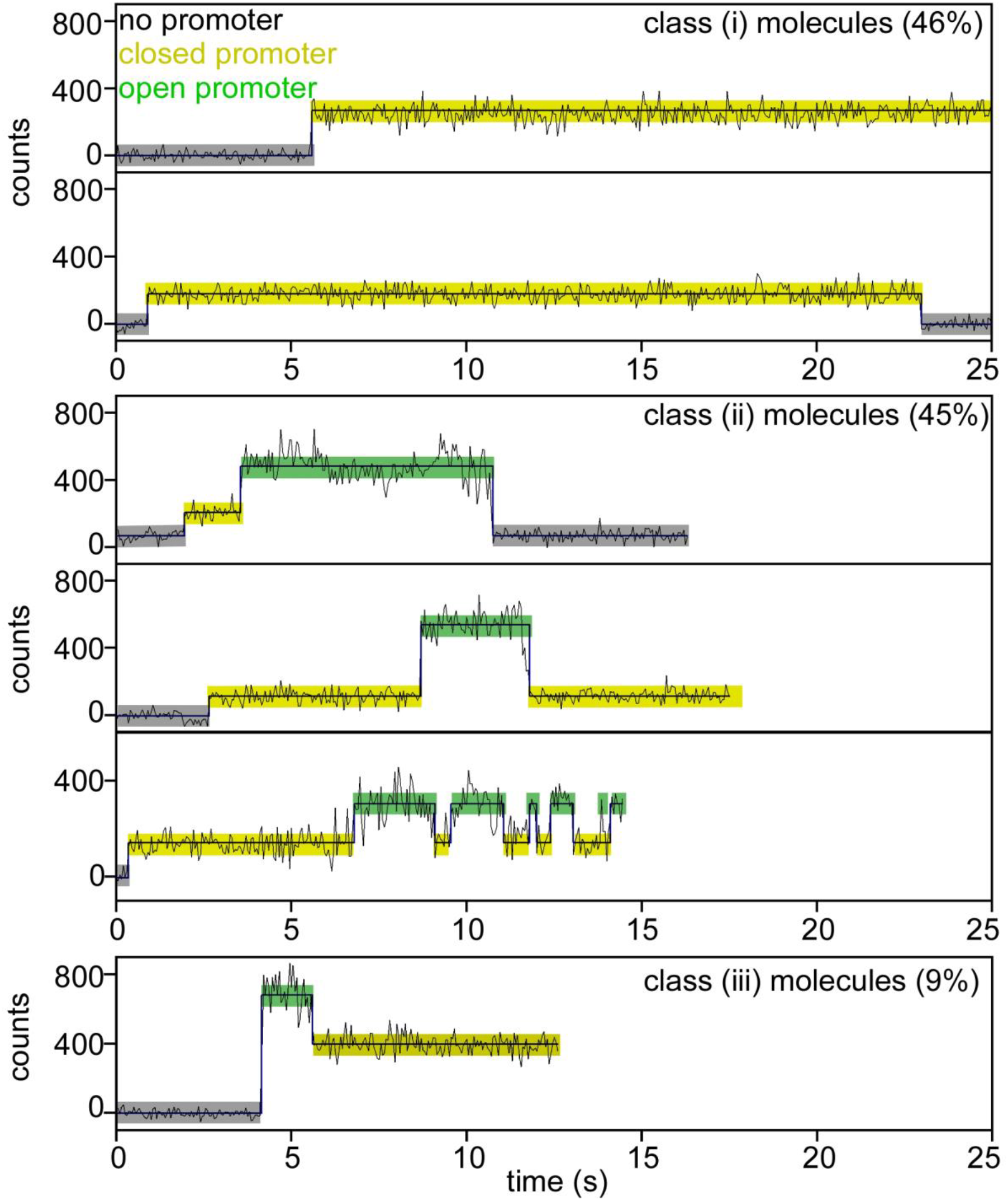
Classification for different time-trajectories of intensity from lacCONS-[+2 Cy3] promoter fragment. Black, raw intensity; dark blue, idealised intensity; Hidden Markov Model (HMM)-assigned states: no promoter (black bars), closed promoter (light yellow bars) and open promoter (green bars). Frame duration: 50-ms; Laser power: 0.60 mW.

**Fig. S4.**
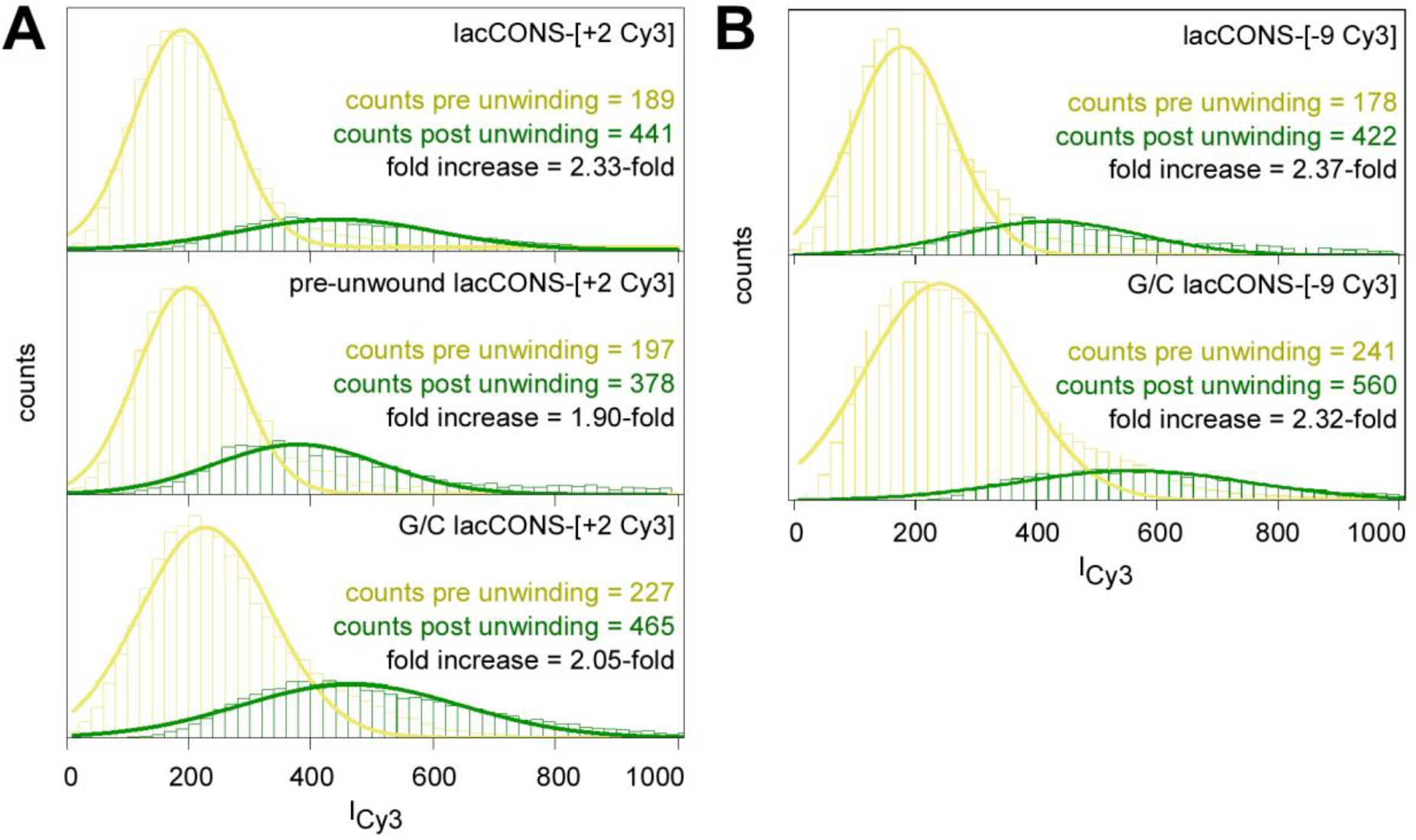
Fluorescence intensities from Cy3-labelled promoter fragments pre-and post-unwinding. HMM-assigned histograms and Gaussian fits of fluorescence intensities, pre-unwinding (yellow) and post-unwinding (green) for Cy3-labelled promoter fragments. Mean intensities for the two states and fold-increase of intensities are shown (*inset*). **A**. Cy3 on +2 position of the non-template strands. **B**. Cy3 on −9 of the template strands.

**Fig. S5.**
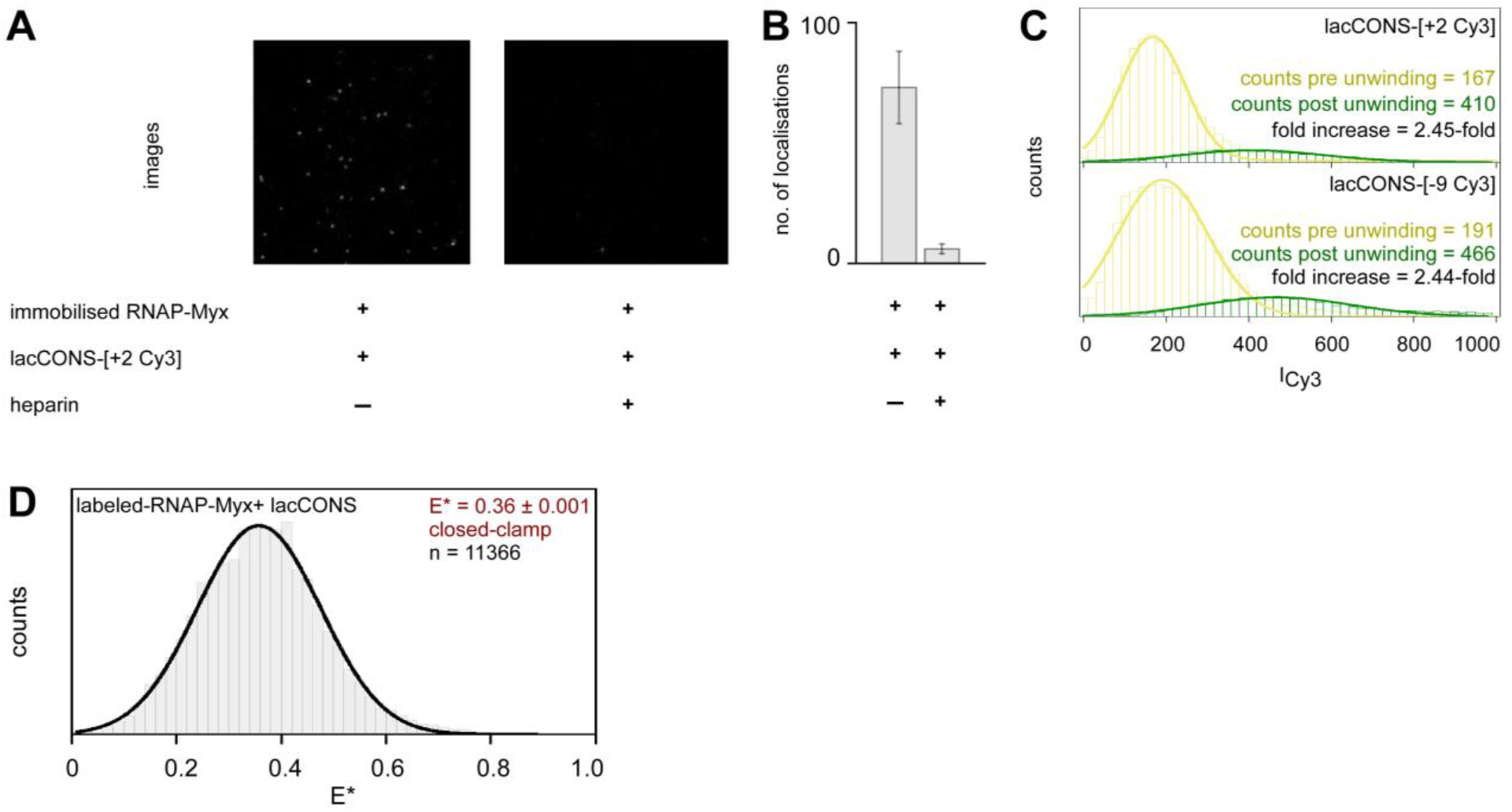
Characterisation of complexes formed between RNAP and promoter fragments in presence of Myxopyronin (Myx). **A**. Raw images showing field of view (24 μm × 24 μm) with immobilised biotin promoter fragment bound to labelled RNAP complexes formed in presence of Myxopyronin (Myx) (*left*), and with remaining immobilised hexahistidine-tagged RNAP holoenzyme complexes bound to Cy3-labelled promoter fragment, formed in presence of Myx, after addition of heparin (*right*). **B**. Mean number of localisations per field of view for single Cy3-labelled promoter fragments bound to immobilised hexahistidine-tagged RNAP holoenzyme, formed in presence of Myx, before and after addition of heparin. Error bars represent the standard deviation from the mean. **C**. HMM-assigned histograms and Gaussian fits of fluorescence intensities corresponding to pre-unwinding (yellow) and post-unwinding (green) states for promoter fragments with Cy3 on +2 position of the non-template strand (*top*) or −9 position of the template strand (*bottom*) of the promoter bubble. Mean intensities for the two states and fold-increase of intensities are shown (*inset*). **D**. Histogram and Gaussian fit of E* showing mean E* values for clamp conformational state for clamp-labelled RNAP molecules bound to immobilised lacCONS-promoter fragments in presence of Myxopyronin.

**Fig. S6.**
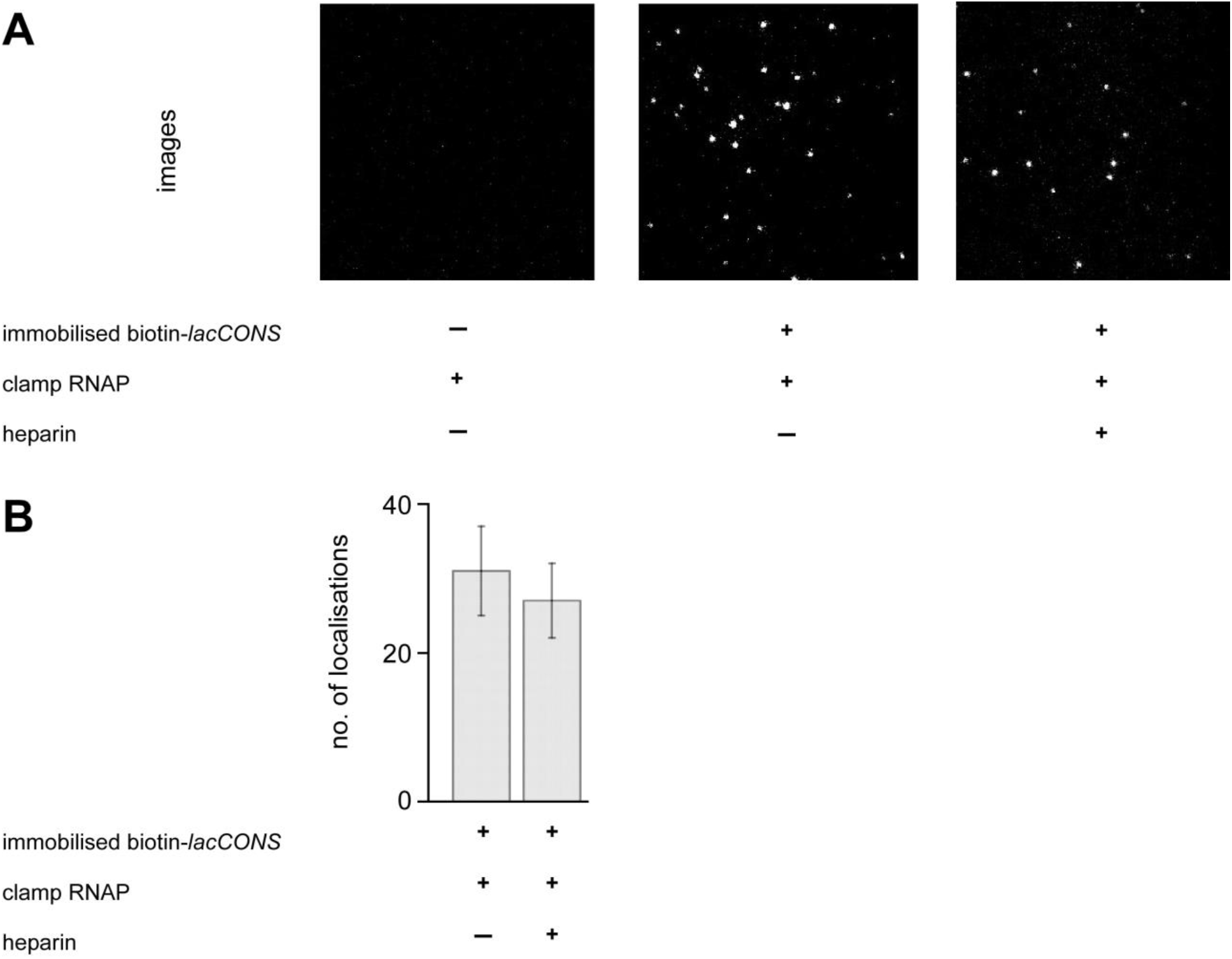
Characterisation of complexes formed between clamp-labelled RNAP and immobilised promoter fragments. **A**. Raw images of the green emission channel showing field of view (24 mm × 24 mm) with immobilised biotin-lacCONS promoter fragments (*left*), with immobilised biotin-lacCONS promoter fragments bound to clamp-labelled RNAP (*middle*), and with remaining immobilised biotin-lacCONS promoter fragments bound to clamp-labelled RNAP, after addition of heparin (*right*). **B**. Mean number of localisations per field of view for single clamp-labelled RNAP molecules with both green (Cy3B) and red (Alexa647) probes, bound to immobilised biotin-lacCONS promoter fragments, before and after addition of heparin. Error bars represent the standard deviation from the mean.

**Fig. S7.**
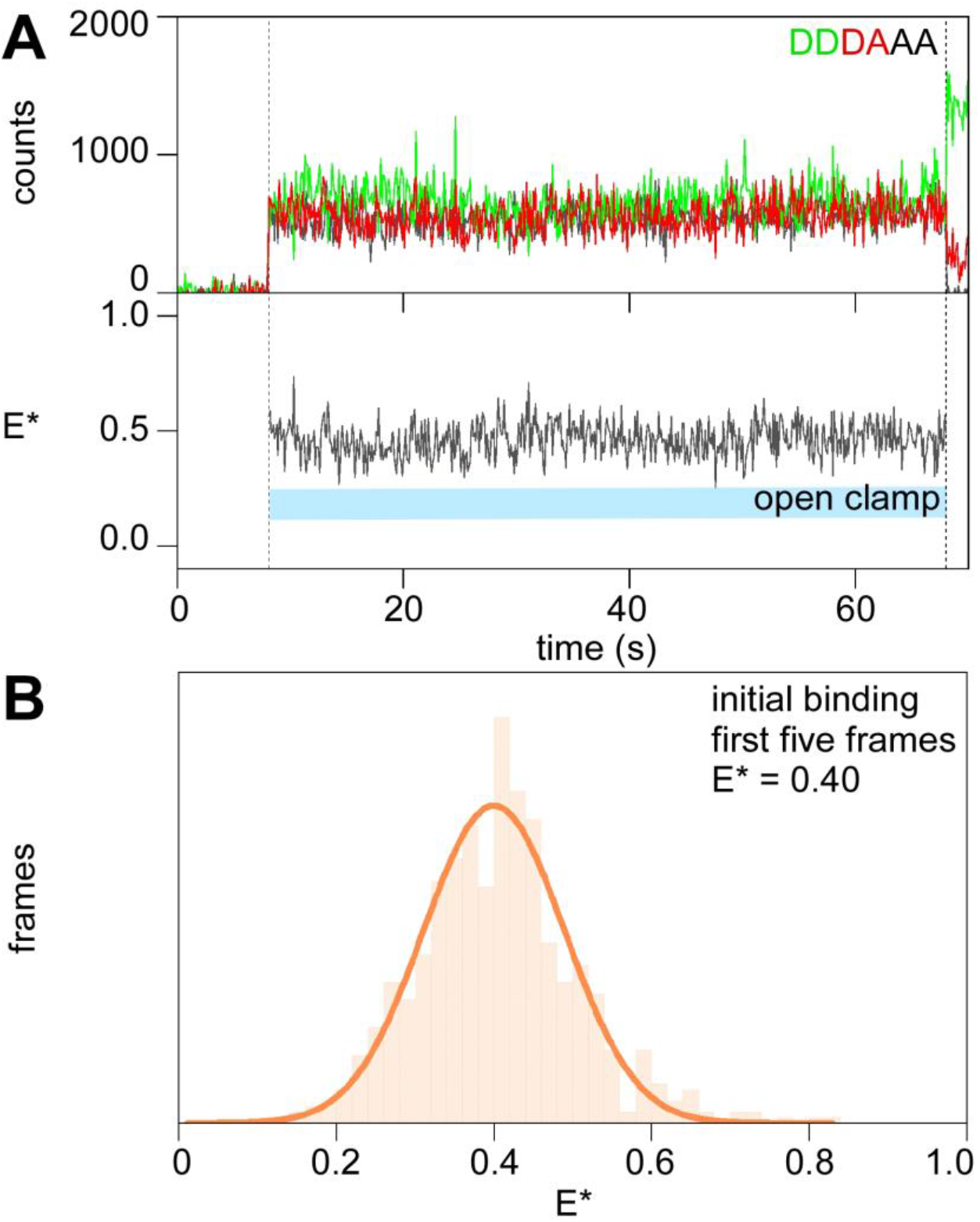
smFRET: initial binding of RNAP to surface immobilised promoter DNA fragments take place via a closed-clamp conformation. **A**. Fluorescence intensity-vs-time trajectories showing simultaneous appearance (at ∼8 s) of donor and acceptor signals from labelled-RNAP bound to promoter (*top*) and FRET efficiency, E* (*bottom*). Range for expected E* values corresponding to an open-clamp conformation is highlighted in cyan. **B**. Histogram and Gaussian fit of E* values for first five frames after binding define the mean E* for initial binding. Frame duration: 100-ms. Laser powers: 200 μW in red and 500 μW in green.

**Fig. S8.**
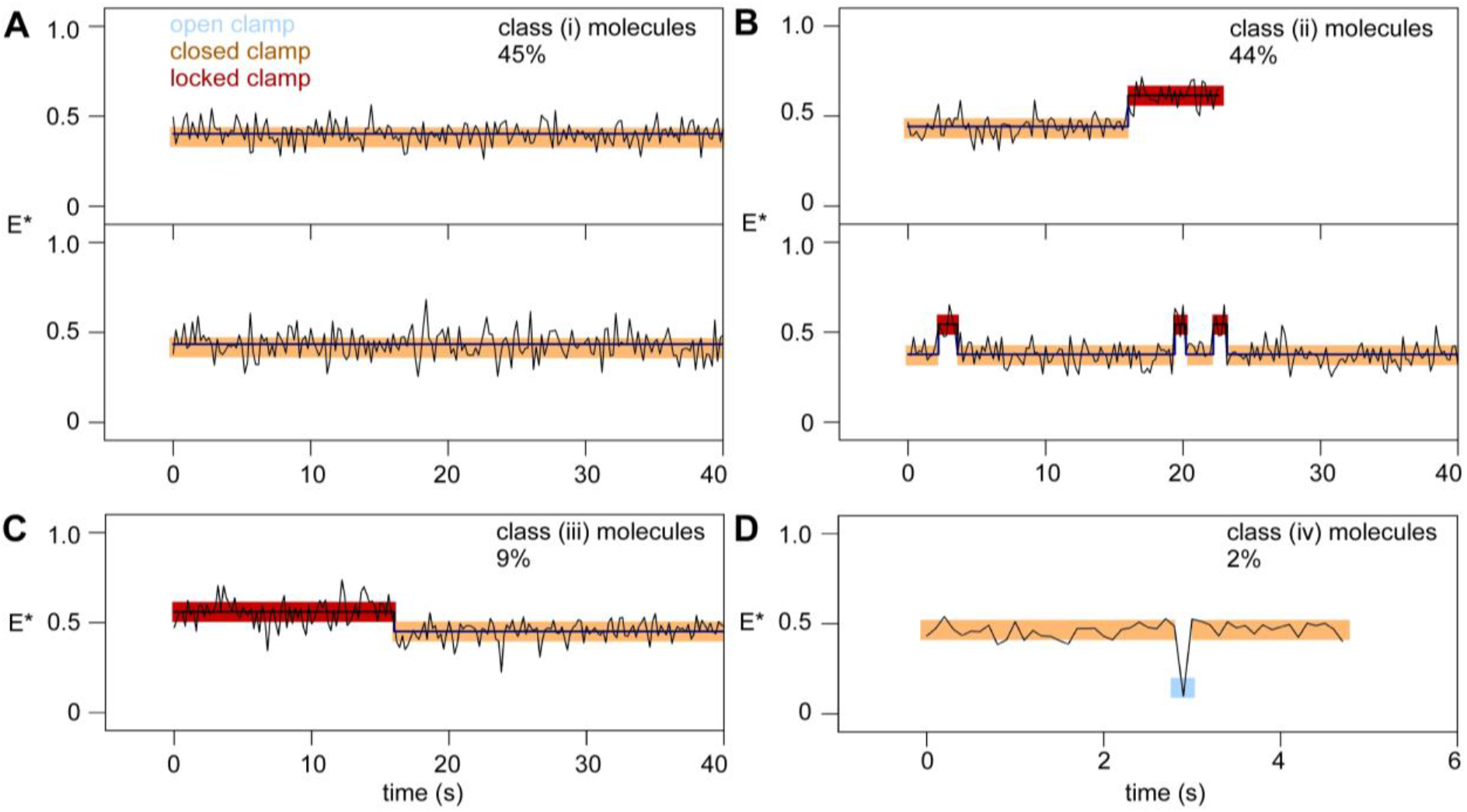
Classification for different time-trajectories of E* of clamp labelled RNAP molecules bound to immobilised lacCONS-promoter fragments in real time. HMM-assigned E* time-trajectories showing closed-clamp states (orange), locked-clamp states (red), open-clamp states (cyan) and interstate transitions (dark-blue).

**Fig. S9.**
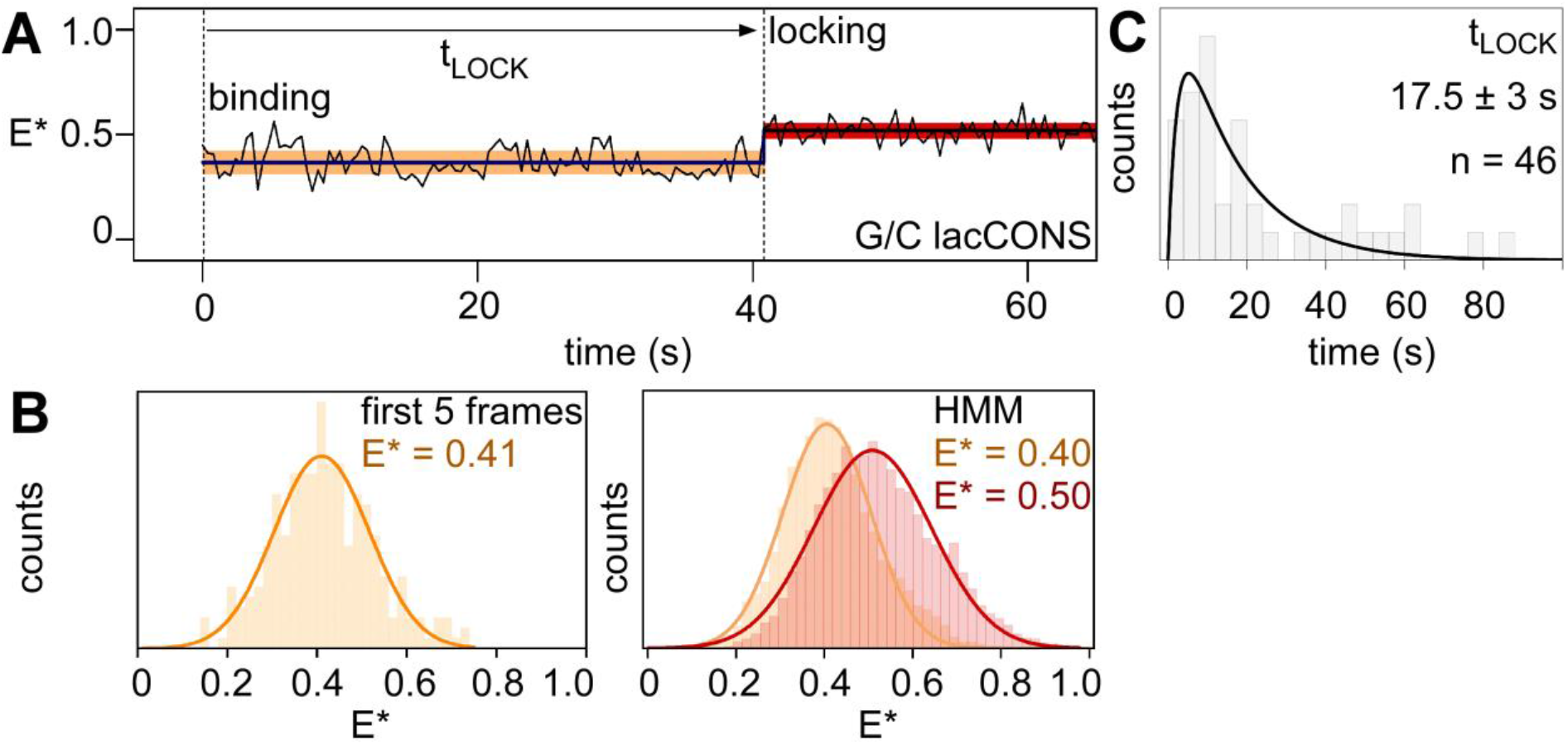
smFRET data showing binding of clamp-labelled RNAP to immobilised biotin-G/C lacCONS promoter fragments. **A**. Time-trajectory of E*, showing HMM-assigned closed-clamp state (orange) and locked-clamp state (red). Frame duration: 400-ms. Laser powers: 50 μW in red and 150 μW in green. **B**. HMM-assigned histograms and Gaussian fits of E* for first five-frames (*left*) and full time-trajectories (*right*). **C**. Dwell-time histograms for time before transition to the locked-clamp state, t_LOCK_.

